# Operationalizing site-level conservation for migratory birds across the Americas’ flyways

**DOI:** 10.64898/2026.08.12.744493

**Authors:** Daniela Linero-Triana, Nathaniel E. Seavy, Santiago Aparicio, Jhan C. Carrillo-Restrepo, Rob Clay, Olivia Crowe, William V. De Luca, River Gates, Victoria Jones, Arne Lesterhuis, Nicole L. Michel, Michael Seager, Maria Gabriela Toscano, Jhonnattan Valdés-Uribe, Mauricio Velásquez, Jorge Velásquez-Tibatá

## Abstract

Conserving migratory birds effectively requires full annual cycle strategies that identify where on-the-ground efforts can have the greatest impact. Here, we present a hemispheric spatial framework to identify priority areas for 112 migratory bird species across the Americas. Building on full annual cycle prioritizations, we defined finer-scale spatial planning units that reflect differences in migratory and congregational behaviors between shorebirds and landbirds. We compiled population data for each planning unit and focal species and applied conservation planning tools to design area-efficient portfolios of sites and landscapes that secure 10% of each species’ population within the Americas’ flyways. The resulting minimum area portfolios include 175 shorebird sites and 80 landbird landscapes optimized to meet the species-specific 10% representation targets across breeding, non-breeding, and passage seasons. We also identified a broader set of complementary solutions, ranked by an importance score, to provide decision-makers with flexible options for strategic resource allocation. This framework provides the scientific foundation for the Americas Flyways Initiative (AFI), which aims to catalyze investment in nature-based solutions and bird-friendly infrastructure to enhance the conservation of migratory birds and strengthen the resilience of the Americas’ flyways by 2050.

## Introduction

Each year, billions of migratory birds undertake extraordinary journeys across continents and oceans, linking regions separated by thousands of kilometers (Runge et al., 2014; Somveille et al., 2015). These large-scale movements enable birds to avoid the harsh temperate zone winters and exploit seasonal resource pulses, enhancing survival and reproductive success throughout their life cycles (Somveille et al., 2015). Beyond its biological significance, migration drives profound spatial and temporal changes in bird communities (Somveille et al., 2013) and contributes substantially to the global flow of biomass, nutrients, and energy (Horns & Şekercioğlu, 2018; Runge et al., 2015). In the Americas, migratory birds travel along major flyways that span breeding areas in temperate North and South America, stopover sites, and non-breeding grounds in tropical biomes (Jahn et al., 2004; Somveille et al., 2013). Because these movements span vast and dynamic areas facing diverse threats, conservation frameworks must be developed at the flyway scale to encompass the full annual cycles of migratory birds (DeLuca et al., 2023; Kirby et al., 2008).

The urgent need for flyway-scale approaches is underscored by the ongoing decline of migratory bird populations. In the past half-century, North America alone has lost nearly 3 billion birds (Rosenberg et al., 2019), and recent assessments indicate that this decline persists (North American Bird Conservation Initiative, 2025; UNEP-WCMC, 2024). Migratory species encounter a wide range of threats throughout their annual cycles, including habitat loss and degradation, movement barriers, collisions, predation, harvest and overexploitation (Kirby et al., 2008; López-Hoffman et al., 2017; Andres et al., 2022; Seavy et al., 2025). These threats are spatially and seasonally variable, and their effects often accumulate or interact synergistically (Buchan et al., 2023; Nemes et al., 2023; Seavy et al., 2025). For instance, climate change is expected to intensify many of these pressures and further alter habitat conditions (Kirby et al., 2008; López-Hoffman et al., 2017). Hence, addressing these challenges at the flyway scale is essential for safeguarding migratory birds and maintaining their broader ecological functions (DeLuca et al., 2023; Guo et al., 2024; Kirby et al., 2008; López-Hoffman et al., 2017).

However, conservation at the flyway level is challenging due to its hemispheric scope and the diversity of stakeholders involved (Horns & Şekercioğlu, 2018; Martin et al., 2007). Nevertheless, flyway-scale conservation presents a unique opportunity to foster multisectoral collaboration and transnational partnerships by positioning migratory birds as ambassadors for international conservation (Horns & Şekercioğlu, 2018; Roca et al., 1996). In practice, these broad frameworks can be implemented effectively for many species by dynamically identifying priority sites where on-the-ground efforts will have the greatest impact. For migratory species, it is crucial to recognize that the outcomes of actions in one location or season often depend on complementary efforts elsewhere in their range (López-Hoffman et al., 2017; Norris et al., 2004; Runge et al., 2014). Therefore, establishing portfolios of priority areas across the Americas flyways is a critical foundation for an effective and integrated hemispheric conservation framework (Hunter et al., 1991; Kirby et al., 2008).

Recent advances offer new opportunities for identifying portfolios of priority sites. For example, DeLuca et al. (2023) developed hemisphere-scale, full annual cycle prioritizations of broad regions with high conservation value for migratory birds breeding in various North American ecosystems. Building on this hemispheric perspective, finer-scale site and landscape identification allows for more targeted local conservation actions. Advances in species abundance modeling (Fink et al., 2023) and the availability of extensive bird count data from conservation programs and crowd-sourced platforms (Sullivan et al., 2009) now enable the use of complementarity-based algorithms to design efficient portfolios of priority sites that achieve conservation goals and maximize biodiversity impact. These portfolios help ensure that priority areas complement each other, thereby maximizing ecological benefits (Kukkala & Moilanen, 2013).

A portfolio of priority places can be utilized by a wide range of organizations and decision makers. Notably, this study will directly inform the Americas Flyways Initiative (AFI), led by the National Audubon Society, BirdLife International, and CAF-development bank of Latin America and the Caribbean. Inspired by a similar initiative launched in Asia in 2021 (East Asian– Australasian Flyway Partnership, 2023), AFI aims to address biodiversity loss and climate change through innovative funding mechanisms supported by both private and public sources. Its primary goal is to enhance the resilience of the Americas’ flyways by protecting, restoring, and managing sites needed to secure at least 10% of focal migratory bird populations at the flyway level by 2050. Therefore, identifying these places is essential for selecting projects that will optimize the effectiveness of AFI’s prospective investment.

In this study, we identify a portfolio of sites and landscapes critical for the conservation of focal migratory birds across the Americas, with the goal of helping secure 10% of their flyway populations. We also develop an importance score to provide a consistent metric for comparing sites across potential portfolio solutions. Finally, we assess the potential for implementing various nature-based solutions (NbS) and bird-friendly infrastructure projects promoted by AFI, based on the species and threats present at priority sites.

## Methods

### Selection of focal species

To identify focal migratory bird species across the Americas’ flyways, we focused on two major ecological groups: shorebirds and landbirds (Figure 1). These groups encompass a wide range of habitat associations, migratory strategies, and life history traits (Cañizares & Reed, 2020; Myers, 1983). Furthermore, we considered both boreal migrants (breeding primarily in North America) and austral migrants (breeding primarily in southern South America) to capture the full extent of latitudinal migration systems in the Western Hemisphere.

**Figure 1.**
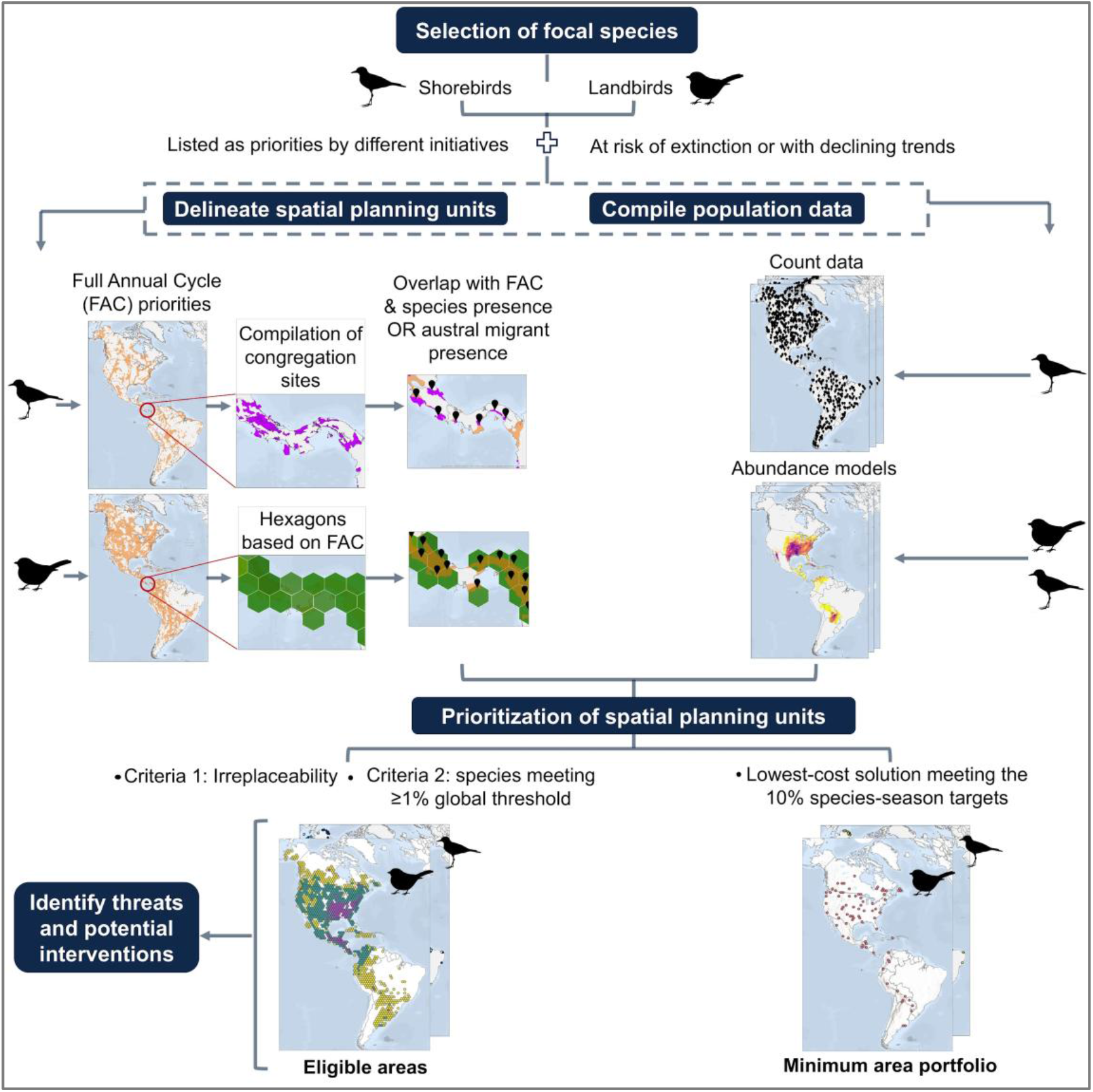
Methodological workflow used to identify priority areas for migratory bird conservation across the Americas’ flyways.

For each ecological group (shorebirds and landbirds), we selected species recognized as being of high conservation concern by regional and hemispheric initiatives. Specifically, for shorebirds, we compiled lists of species from the Atlantic, Midcontinental, and Pacific Shorebird Conservation Partnerships (Atlantic Flyway Shorebird Initiative, 2015; Midcontinent Shorebird Conservation Initiative (MSCI), 2025; Senner et al., 2016), as well as from Birds Caribbean, Alianza Pastizal (Di Giacomo & Parera, 2008), and the Western Hemisphere Shorebird Reserve Network (WHSRN; Western Hemisphere Shorebird Reserve Network, 2020). For landbirds, we included species listed by Alianza Pastizal, Birds Caribbean, the U.S. Fish and Wildlife Service’s Birds of Conservation Concern (U.S. Fish and Wildlife Service, 2021), Partners in Flight (Rosenberg et al., 2016), and the Road to Recovery initiative (Road to Recovery, 2023). This yielded an initial list of 56 shorebird and 258 landbird species of conservation concern.

We applied a second filter according to conservation vulnerability and retained only species classified as at risk of extinction or with declining population trends, based on the IUCN Red List and other conservation assessments (Road to Recovery, 2023; Rosenberg et al., 2016; U.S. Fish and Wildlife Service, 2021). We then excluded species with extremely rare or localized distributions, large year-round resident populations, short-distance migration, limited monitoring feasibility or insufficient data, based on the authors’ expert judgment. The final selection resulted in 51 shorebird and 61 landbird species (Table S1).

### Definition of spatial planning units

We implemented a hierarchical approach to delineate spatial planning units as potential priority areas, identifying relevant places at local and landscape scales while situating them within a hemispheric Full Annual Cycle (FAC) context (DeLuca et al., 2023).

First, we used the FAC prioritizations developed by DeLuca et al. (2023) to identify broad regions of importance for migratory bird populations and connectivity across the Western Hemisphere (Figure 1). These regions served as a spatial framework within which finer-scale planning units were delineated for our focal species. DeLuca et al.’s FAC prioritizations integrate data from tracking, banding, and migratory connectivity to identify areas of high conservation value for birds breeding in six major North American ecosystems. The species selected for each ecosystem were intended to represent habitat-specific bird communities in North America (DeLuca et al., 2023) and therefore do not fully overlap with the focal species compiled for this study. Moreover, the FAC prioritizations do not explicitly consider austral migrants. Despite these limitations, we used the FAC layers as a hemispheric contextual framework within which finer-scale priorities for our focal boreal and austral migrant species could be identified.

Accordingly, for our focal shorebirds, we defined broad regions of high conservation importance using the top 30% of FAC prioritization scores for migratory waterbirds. We selected this threshold as an inclusive yet selective cutoff, consistent with its use in the original application (DeLuca et al. 2023). The FAC waterbird prioritization included 54% of our focal shorebird species. For landbirds, we created an integrated FAC layer by combining five North American ecosystem-based prioritizations (boreal forests, eastern forests, western forests, grasslands, and arid lands). For each ecosystem, we retained the top 30% of priority values and then calculated the maximum value across all five layers, producing a single integrated surface that defined the broad region of importance for landbirds. This integrated layer included 38% of our focal landbird species.

Within these broad FAC regions, we delineated spatial planning units (Figure 1) using different approaches for shorebirds and landbirds to reflect differences in their migratory and congregational behaviors. Shorebirds often concentrate in large numbers at regular stopover sites during migration and in the stationary non-breeding season (Cañizares & Reed, 2020; Myers, 1983), which has enabled multiple established conservation programs to identify key congregation sites. To incorporate this information, we compiled spatial polygons of sites identified as important by the Atlantic, Midcontinental, and Pacific Shorebird Conservation Partnerships (Atlantic Flyway Shorebird Initiative, 2016; Senner et al., 2016), Important Bird and Biodiversity Areas (IBAs, a subset of Key Biodiversity Areas; Donald et al., 2019; BirdLife International, 2022), Important Shorebird Sites (Lesterhuis et al., 2022), Ramsar Wetlands of International Importance (Ramsar Convention on Wetlands, 2022), and the Western Hemisphere Shorebird Reserve Network (WHSRN, 2023). Only sites overlapping waterbird FAC priority regions and with occurrences of focal shorebirds were retained (see “Population data acquisition for focal species”).

Two challenges arose during this process: (1) some sites appeared in multiple conservation programs with different polygon geometries, and (2) some sites were only represented as points. We manually merged overlapping entries with the same site name into a single polygon. For the remaining point-only sites, we applied a 10 km buffer to approximate the median polygon area of the compiled dataset.

For landbirds, we used a different approach due to their limited availability of site-based flyways initiatives comparable to those for shorebirds and their tendency to migrate across broad fronts (Guo et al., 2024; Runge et al., 2015). Given this migratory behavior, conservation efforts for these species are typically implemented at the landscape scale (Guo et al., 2024; Runge et al., 2015). Hence, we divided the landbird FAC integrated prioritization into ∼20,000 km² hexagons, which we termed ’landscapes’ and selected those with more than 25% of their area within the FAC priority areas. As with shorebirds, we retained landscapes with reported occurrences of focal landbirds (see “Population data acquisition for focal species”).

Finally, because DeLuca et al.’s FAC prioritizations did not include austral migrants, we ensured their representation by retaining any spatial planning unit with occurrences of focal austral species, even if it fell outside or only minimally overlapped the FAC broad priority regions.

### Population data acquisition for focal species

For shorebirds, we gathered count data to capture patterns of localized abundance at key sites, compiling records from eBird (eBird, 2023), the Global Biodiversity Information Facility (GBIF), BirdLife’s World Bird and Biodiversity Database (WBDB), Manomet’s important shorebird sites database (Lesterhuis et al., 2022), and published literature (Supplementary File 1). We included occurrences up to December 31, 2022, within defined spatial planning units. We cleaned eBird and GBIF records by removing duplicates and retaining only those with area, stationary, traveling, or International Shorebird Survey protocols. Each count was assigned to a migratory season based on its geographic location relative to the species’ distribution ranges (BirdLife International and Handbook of the Birds of the World, 2021).

For landbirds, and to complement shorebird counts, we used weekly 3 km-resolution proportion-of-population rasters from the eBird Status and Trends models (eBirdST; Fink et al., 2023). For species with available eBirdST models, seasonal population percentages were calculated using eBird migration date ranges. For species without eBirdST models (*Sporophila cinnamomea*, *Sporophila palustris*, and *Sporophila ruficollis*), we first derived analogous weekly 3 km proportion-of-population surfaces from GBIF records and then inferred seasonal timing from distribution maps (BirdLife International and Handbook of the Birds of the World, 2021). For both data sources, breeding and non-breeding population estimates were calculated as the maximum pixel value during the respective season. For the pre- and post-breeding periods (passage), we assumed conservative two-week turnover intervals (DeLuca et al., 2021) and summed population sizes every two weeks for each pixel. We aggregated population percentages across all pixels within each unit and species. Because shorebird counts are reported as population sizes rather than percentages, we converted eBird-derived percentages into population counts by multiplying them by available global population estimates (Andres et al., 2012; Rosenberg et al., 2016; Wetlands International, 2023). Finally, to focus on migratory populations, we excluded estimates for shorebirds and landbirds within year-round resident ranges (BirdLife International and Handbook of the Birds of the World, 2021).

### Portfolio identification and prioritization of spatial units

We identified the spatial planning units required to secure 10% of the population of all focal species using Marxan 2.43 (Ball et al., 2009), which optimizes combinations of units that meet representation targets at the lowest total cost.The 10% representation targets were defined per species and per season (breeding, non-breeding, and passage) based on flyway-scale population estimates. For each species and season, flyway population size was calculated as the sum of the maximum population values across all spatial planning units that met the filtering criteria. For shorebirds, these seasonal values incorporated both maximum count data and eBirdST–derived population sizes. For landbirds, seasonal values were derived from eBirdST-based population percentages. We used the area of each unit as a proxy for cost (terrestrial area for landbird units). Marxan was run 100 times, and the solution with the lowest total cost was selected as the minimum-area portfolio (Figure 1). We did not consider the representation of focal species in existing protected areas, as protected areas and other managed areas could still be eligible for AFI funding, if they implement actions to minimize threats.

Because multiple alternative solutions could meet the same targets, albeit at a higher overall cost, we quantified the relative importance of all planning units by calculating an importance score based on two prioritization criteria (PC; Figure 2). PC1 measured the irreplaceability of each unit in achieving the 10% representation target; values near 100 indicate essential areas, whereas lower values suggest many possible replacement alternatives. A value of zero indicates that the unit does not contribute to meeting the representation targets. PC2 was defined as the number of species for which the unit held at least 1% of the global population, a commonly used threshold in multiple conservation programs (IUCN, 2016).

**Figure 2.**
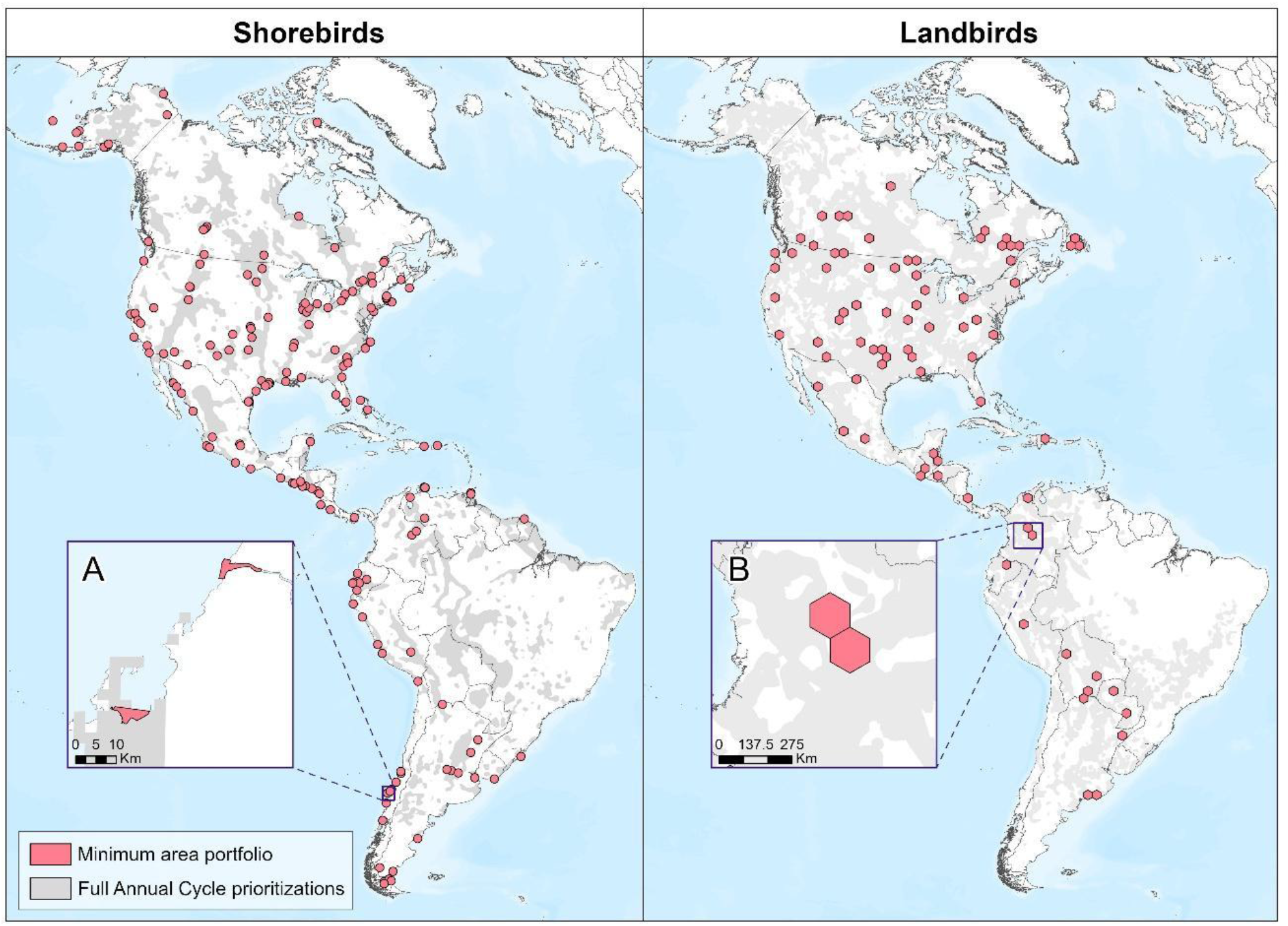
Minimum area portfolio for shorebirds and landbirds, representing the set of spatial planning units that achieved 10% of seasonal population targets at the lowest total cost. Shorebird units are shown as points at the hemispheric scale for clarity. (A) Example of shorebird units along the Chilean coast. (B) Example of landbird units in midwestern Colombia.

PC1 and PC2 were classified into three categories: for PC1, low (<50), medium (50–89), and high (90–100); for PC2, low (below the median), medium (median to the 90th percentile), and high (above the 90th percentile). Units with scores above zero in at least one PC are hereafter referred to as eligible for AFI investments. Units scoring zero in both PCs were excluded from subsequent analyses, although they may still be considered candidates for investment under specific conditions (Figure S1). The final importance score for each eligible unit corresponded to the highest score between PC1 and PC2.

### Assessment of threats and conservation opportunities

We used published spatial datasets to assess potential threats to focal migratory birds within planning units and to identify opportunities for implementing NbS or minimizing the impacts of existing infrastructure projects on birds (Figure 1). We quantified the extent of croplands, urban infrastructure, rangelands, and roads in each unit. Rangelands were treated as potential threats due to livestock grazing; however, to avoid overestimation, particularly in tundra ecosystems, we excluded areas that did not overlap with the livestock management layer of Seavy et al. (2025). All other threats were assessed as presence/absence because available hemispheric datasets were either too coarse in resolution or reported only point locations, preventing reliable estimation of affected area.

To ensure a species-specific assessment, we identified relevant threats for each focal species by reviewing the IUCN Red List, Billerma et al. (2022) and Smith et al. (2022). Only threats documented to affect the focal species present within each planning unit were retained. Based on the resulting threats, we determined potential NbS or infrastructure-related intervention opportunities for each unit (Table 1).

**Table 1.** Summary of spatial data sources used to evaluate potential threats, potential nature-based solutions (NbS) or infrastructure-based interventions across planning units.

| General IUCN-CMS threats classification<sup>1</sup> | Specific IUCN-CMS threats classification | Evaluation | Potential interventions | Source |
| --- | --- | --- | --- | --- |
| 1 Residential and commercial development | Built-up areas | Area and presence | Bird-friendly urban infrastructure | Zanaga et al. (2022) |
|  | Light pollution | Presence |  | Seavy et al. (2025) |
| 2.1 Annual and perennial non-timber crops | Croplands | Area and presence | Sustainable management practices and ecological restoration | Zanaga et al. (2022) |
| 2.3 Livestock farming & ranching | Rangelands | Area and presence |  | Seavy et al. (2025); Zanaga et al. (2022) |
| 2.1 Annual and perennial non-timber crops | Susceptibility to future agricultural development | Presence | Habitat protection | Oakleaf et al. (2019) |
| 3.3 Renewable energy | Solar farms | Presence | Best management practices for birds and clean energy | Global Energy Monitor (2023) |
|  | Wind turbines | Presence |  | Seavy et al. (2025) |
|  | Susceptibility to future solar and wind renewable energy development | Presence |  | Oakleaf et al. (2019) |
| 4.1. Roads and railroads | Roads | Area and presence | Bird-friendly linear infrastructure | Meijer et al. (2018) |
| 4.2. Utility and service lines | Powerlines | Presence |  | Seavy et al. (2025) |
|  | Oil and gas pipelines | Presence |  | Seavy et al. (2025) |
| 7.2. Dams and water management/use | Surface water management | Presence | Management for improved water access, sanitation, and irrigation | Seavy et al. (2025) |
| 9.1. Domestic and urban waste water<br>9.3. Agricultural and forestry effluents | Low water quality | Presence |  | Seavy et al. (2025) |
| 11.1 Habitat shifting & alteration<br>11.4 Storms & flooding | Coastal disturbance | Presence | Coastal management for resilience | Seavy et al. (2025) |
|  | Coastal modification | Presence |  | Seavy et al. (2025) |
|  | Sea level rise | Presence |  | Seavy et al. (2025) |

<sup>1</sup> Based on Salafsky et al. (2025)

## Results

The minimum-area portfolio, optimized to meet species–season 10% representation targets while minimizing total area, comprised 175 spatial planning units for shorebirds and 80 for landbirds (Figure 2), covering 124,342 km² and 1,558,846 km², respectively. The shorebird and landbird portfolios overlapped by 1,206 km². At the national level, the United States contained the largest share of planning units for both groups, accounting for 45% of shorebird units and 44% of the total landbird portfolio area. Across Latin America and the Caribbean, Chile (8%), Mexico (7.4%), and Argentina (5.7%) contributed the highest proportions of shorebird planning units, whereas Mexico (5%), Bolivia (4.4%), and Argentina (4.1%) accounted for the largest shares of landbird portfolio area.

Eligible areas, which include the minimum-area portfolio and represents a broader set of complementary solutions, comprised 603 planning units for shorebirds and 1,040 for landbirds (Figure 3). Relative to the minimum-area portfolio, this expanded set of eligible areas provided substantially greater geographic coverage across the Americas, encompassing 979,791 km² for shorebirds and 20,264,994 km² for landbirds. The eligible shorebird and landbird areas overlapped by 350,031 km². As in the minimum-area portfolio, the United States contained the largest share of planning units for both groups, accounting for 43% of shorebird units and 36% of the total landbird eligible area. Across Latin America and the Caribbean, the highest concentrations of shorebird planning units occurred in Mexico (8%), Argentina (7.3%), and Chile (6.6%), whereas Mexico (7.5%), Brazil (6.2%), and Argentina (5.1%) contributed the largest shares of landbird eligible area.

**Figure 3.**
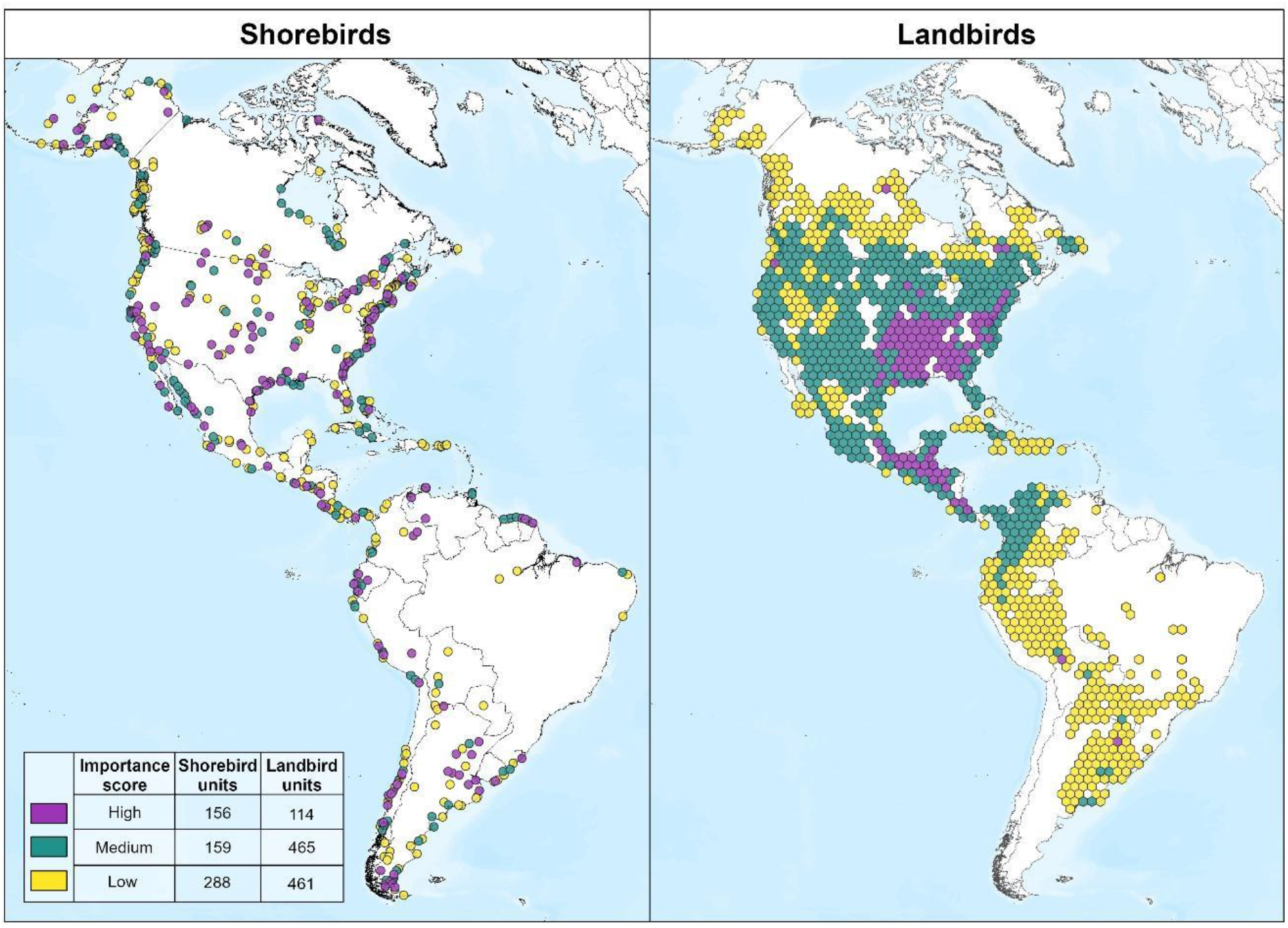
Importance scores for all eligible spatial planning units for shorebirds and landbirds. The accompanying table indicates the number of units within each importance category, based on the highest value between the irreplaceability criterion (PC1) and the number of species reaching the 1% global population threshold criterion (PC2).

Among shorebird planning units, all eligible areas were already recognized under existing conservation designations. Specifically, 71% of eligible units and 80% of those in the minimum area portfolio were designated as IBAs (Table 2). In addition, 51% (n = 306) of eligible units held more than one conservation designation, and 17% (n = 101) were recognized by more than three programs, underscoring their broad conservation relevance.

**Table 2.** Number of shorebird spatial planning units in the minimum area portfolio and in the full set of eligible areas that are designated under existing conservation programs. Many units have more than one formal designation.

| Designation Category | Minimum Area Portfolio Units (No.) | Eligible Planning Units (No.) |
| --- | --- | --- |
| Important Bird and Key Biodiversity Areas (IBAs/KBAs) | 140 | 430 |
| Important Shorebird Sites (Manomet) | 49 | 239 |
| Ramsar Wetlands of International Importance | 39 | 120 |
| Western Hemisphere Shorebird Reserved Network Sites (WHSRN) | 34 | 101 |
| Pacific Flyway Shorebird Initiative (PSCI) | 24 | 92 |
| Atlantic Flyway Shorebird Initiative (AFSI) | 22 | 125 |

Eligible planning units for both shorebirds and landbirds were assigned an importance score corresponding to the highest value between PC1 (irreplaceability) and PC2 (number of species meeting ≥ 1% of the global population). On average, shorebird units had higher irreplaceability values (mean ± SD = 56 ± 40.7) than landbird units (15 ± 19.3; Figure 4A). In contrast, landbird units generally supported more species surpassing the 1 % global population threshold (7 ± 7) than shorebird units (2 ± 2; Figure 4B). When comparing overall focal species richness, differences between groups were smaller, with shorebird units hosting an average of 22 ± 10 species, while landbird units contained 17 ± 12 species (Figure 4C).

**Figure 4.**
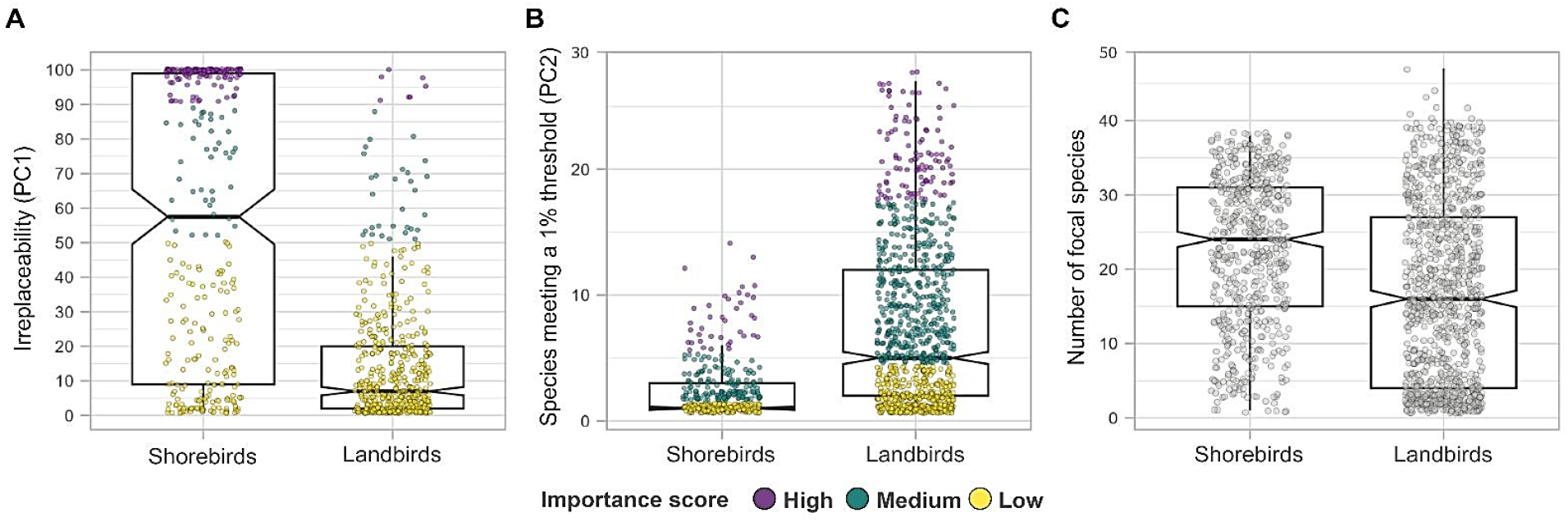
Distribution of prioritization criteria (PC) and focal species richness across eligible spatial planning units for shorebirds and landbirds. (A) PC1: irreplaceability values; (B) PC2: number of species meeting ≥ 1% of their global population; and (C) number of focal species per unit.

When examining species-level representation, all 51 focal shorebird species met the 10% population target during the breeding and non-breeding seasons within the set of eligible areas (Figure 5). During migration, only one of the 35 species with a defined passage range based on distribution maps, the Snowy Plover (*Charadrius nivosus*), fell below the threshold, achieving 8% representation. In the minimum area portfolio, which includes fewer planning units, a slightly higher number of species did not meet seasonal targets (Figure 5).

**Figure 5.**
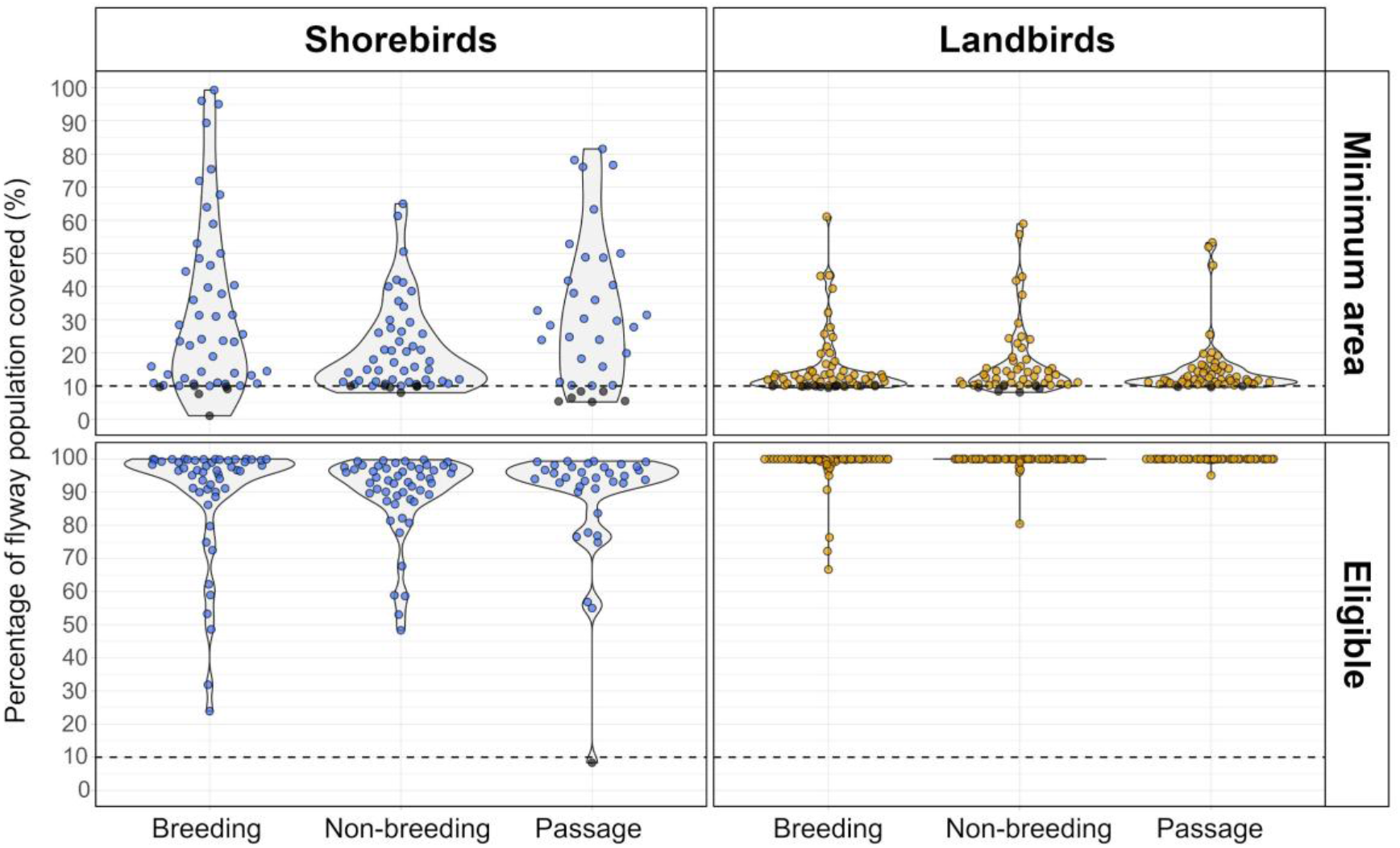
Distribution of seasonal flyway populations represented in the minimum area portfolio and the eligible planning units. Circles correspond to individual shorebird (blue) and landbird (orange) species, with black circles indicating species below the 10% population representation target.

For landbirds, the set eligible areas achieved all species–season targets for focal species that had modeled populations within the spatial planning units. However, several species had no modeled abundance across the planning units in certain seasons and therefore no representation targets. Species without abundance data during the non-breeding and passage seasons included the Chestnut Seedeater (*Sporophila cinnamomea*), Dark-throated Seedeater (*Sporophila ruficollis*), Marsh Seedeater (*Sporophila palustris*), Colima Warbler (*Leiothlypis crissalis*), and Elf Owl (*Micrathene whitneyi*). Additionally, three species lacked non-breeding abundance data: the Chimney Swift (*Chaetura pelagica*), Connecticut Warbler (*Oporornis agilis*), and Flammulated Owl (*Psiloscops flammeolus*). Together, species without abundance data and therefore without representation targets are 7% of all focal species–season combinations. In the minimum-area portfolio for landbirds, targets were not met for a small subset of species, particularly during the breeding season (n = 11; Figure 5).

In relation to existing conservation coverage, we examined how the identified spatial planning units intersect with protected areas, as AFI investments may occur both within and outside their boundaries. Both the minimum area portfolio and the full set of eligible areas for shorebirds showed greater representation within protected areas, particularly under IUCN Category V (Habitat or Species Management Area), compared to landbirds (Table 3).

**Table 3.** Area percentage of spatial planning units under different IUCN protection categories for the minimum area portfolio and the full set of eligible areas. Larger percentages for each conservation planning portfolio are underscored.

| IUCN Protected area category | Shorebirds |  | Landbirds |  |
| --- | --- | --- | --- | --- |
|  | Minimum area portfolio coverage (%) | Eligible planning units coverage (%) | Minimum area portfolio coverage (%) | Eligible planning units coverage (%) |
| I a. Strict Nature Reserve | 0.20 | 0.71 | 0.08 | 0.31 |
| I b. Wilderness Area | 0.02 | 0.13 | 0.34 | 0.51 |
| II. National Park | 0.23 | 1.26 | 1.11 | 0.79 |
| III. Natural Monument | 0.04 | 4.61 | 0.27 | 0.25 |
| V. Habitat or Species | <u>65.61</u> | <u>15.50</u> | 0.89 | 1.07 |

| Management Area |  |  |  |  |
| --- | --- | --- | --- | --- |
| V. Protected Landscape/Sea scape | 4.32 | 3.58 | <u>1.90</u> | 1.66 |
| VI. Protected Area with sustainable use | 1.89 | 5.14 | 0.38 | 0.87 |
| No category | 3.59 | 4.32 | 1.47 | <u>2.00</u> |

Finally, when evaluating threats in terms of spatial extent within planning units, conversion of natural habitats to rangelands and croplands were the most extensive pressures across all units (Figure 6). These land uses, represent opportunities for implementing NbS that reduce threats and enhance habitat quality, such as sustainable management practices and ecological restoration. When threats were evaluated based on their occurrence rather than area, croplands and rangelands remained among the most recurrent pressures; however, built-up areas affected the largest percentage of planning units (Figure 7), highlighting opportunities to improve the bird-friendliness of urban environments.

**Figure 6.**
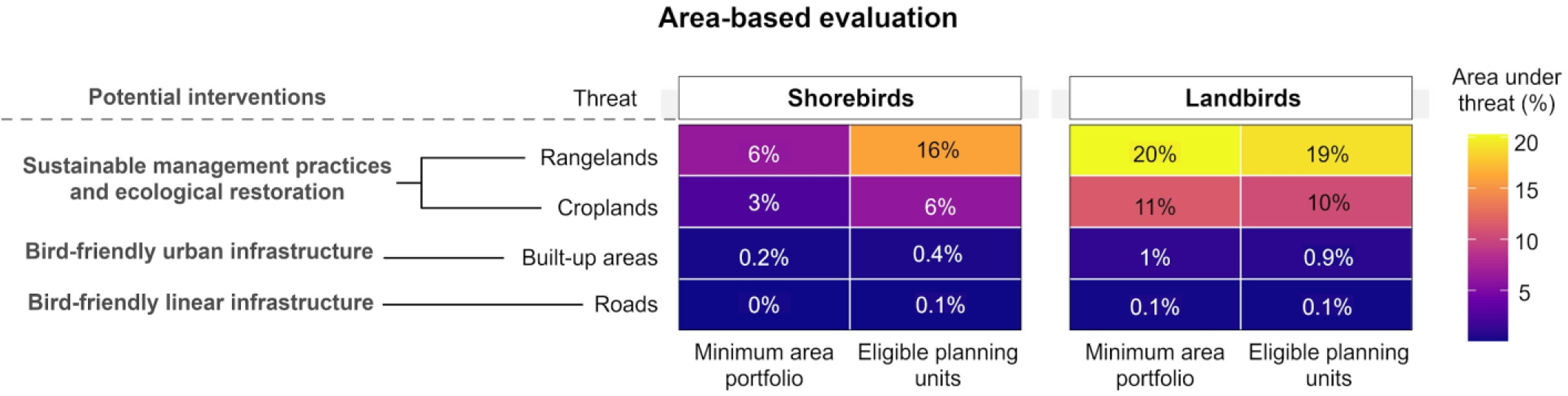
Percentage area coverage of the minimum area portfolio and eligible planning units impacted by different threats, along with potential nature-based or infrastructure-based solutions. Only threats affecting focal species within each planning unit were considered.

**Figure 7.**
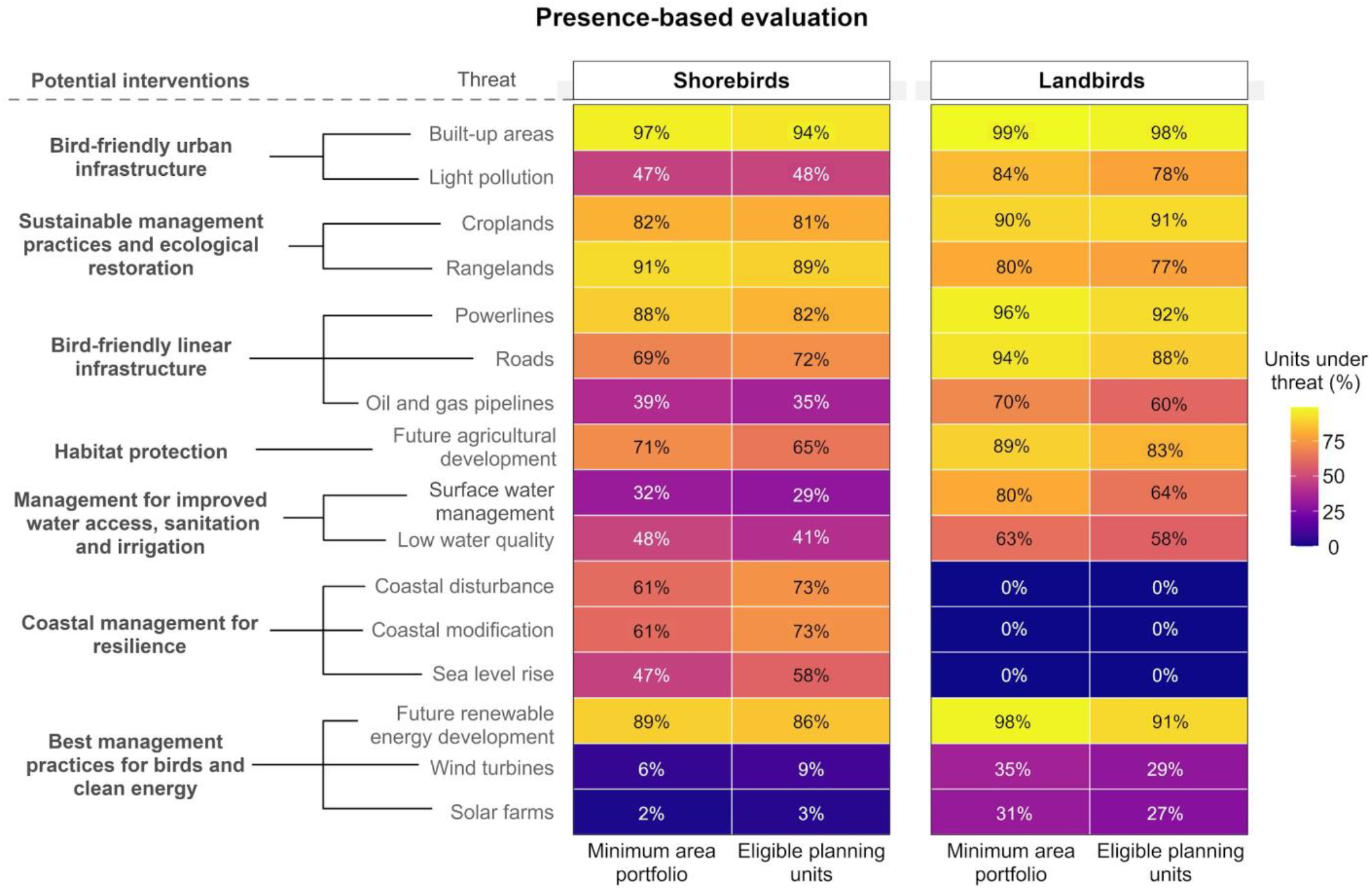
Percentage of planning units in the minimum area portfolio and full set of eligible areas impacted by different threats, along with potential nature-based or infrastructure-based solutions. Only threats affecting focal species within each planning unit were considered.

## Discussion

Historically, migratory bird conservation in the Western Hemisphere has focused on breeding-season actions in North America and on single species facing severe declines (American Bird Conservancy, 2006; SARA, 2002). Yet effective conservation requires identifying priority places that sustain the populations of multiple species throughout their full annual cycles (DeLuca et al., 2023; Kirby et al., 2008). In this study, we delineate sets of spatial planning units that are critical for focal shorebirds and landbirds across the Americas. Building on hemispheric FAC prioritizations (DeLuca et al., 2023), we identified finer-scale key areas comprising 603 units for shorebirds and 1,040 for landbirds, of which 30% and 8%, respectively, form the most area-efficient portfolios. These portfolios were optimized to meet species-specific 10% representation targets across breeding, non-breeding, and passage seasons. By reporting both minimum area portfolios and a broader set of complementary eligible solutions with an importance score, we provide investors, planners, and implementers with actionable options to allocate resources strategically while balancing feasibility and co-benefits for nature and people (Ban et al., 2013; DeLuca et al., 2023; Schwartz et al., 2018).

For shorebirds, our approach leveraged extensive data from multiple established programs that identify sites of biodiversity importance, including IBAs/KBAs and the WHSRN. These programs have long guided governments, NGOs, and donors in prioritizing conservation actions and promoting site conservation through voluntary commitments by responsible entities (Waliczky et al., 2019; WHSRN, 2019). Additionally, these programs document critical congregation sites essential for migratory success (Cañizares & Reed, 2020). Accordingly, several well-known sites, such as Ensenada de Pabellones (Mexico) and Lagoa do Peixe National Park (Brazil), recognized as IBAs, WHSRN sites, and Ramsar wetlands, are among the many included within the minimum-area portfolio. Incorporating existing designations into our analysis ensured that expert knowledge and locally derived information complemented crowd-sourced observations and modeled abundance data (IUCN, 2016).

For landbirds, our prioritization results also align with prior work. Northern Central America has been identified as a focal region for Neotropical terrestrial migrants during the passage and non-breeding seasons, and as a crucial area for conservation investment (Morales et al., 2024; González et al., 2023; Wilson et al., 2019, 2022). Consistent with these findings, our minimum-area portfolio concentrates prioritized units in this region. Likewise, our prioritized landscapes near the Pacific coast of Washington and Oregon overlap areas recognized for supporting large numbers of hundreds of Nearctic-Neotropical migrant species during both spring and fall (Lin et al., 2020a). In South America, our minimum area solution also includes southeastern Bolivia, a region previously highlighted as critical for 117 Neotropical migrants (Schuster et al., 2019).

Across both the shorebird and landbird portfolios, overlap with existing protected areas is limited. For shorebirds, the only notable exception is that more than half of the minimum area portfolio overlaps with IUCN Category V (Habitat or Species Management Areas). Other studies have reported similar protection gaps across migratory cycles (Runge et al., 2015). These findings underscore the importance of strengthening management within existing protected areas and expanding conservation efforts beyond their boundaries, particularly in human-dominated landscapes (Guo et al., 2024; Schuster et al., 2019). NbS and bird-friendly infrastructure are key complementary approaches to protected areas to safeguard, manage, and restore habitats while reducing impacts of urban development and infrastructure (Snep et al., 2016).

Our analysis shows that ranching and agriculture are the most extensive and recurrent threats within priority spatial planning units. These landscapes create opportunities to scale up NbS interventions with documented benefits for birds, such as habitat restoration on private lands (Dertien & Baldwin, 2022; Gonzalez et al., 2023; Michel et al., 2020), silviculture and agroforestry practices (Dertien & Baldwin, 2022; McDermott & Rodewald, 2014), and the creation of temporary wetlands in agricultural matrices (Golet et al., 2018). At the same time, built-up areas, light pollution, power lines, and roads are prevalent in most eligible areas. The global assessment of Simkins et al. (2023) similarly reports widespread infrastructure within KBAs, alongside substantial risk of future development, particularly in the Global South. Targeted interventions to reduce habitat loss, collisions, disturbance and other associated impacts are therefore essential. In the United States alone, buildings, communication towers, power lines, and vehicles cause the deaths of hundreds of millions of birds annually (Loss et al., 2015; Loss, Will, & Marra, 2014; Loss, Will, Loss, et al., 2014). Bird-friendly measures applied to existing infrastructure such as the installation of collision-reduction technologies within areas prioritized in this study could help reduce mortality, particularly where seasonal populations are most concentrated.

The Americas Flyways Initiative (AFI) is well-positioned to turn the spatial priorities of this study into action. It seeks to catalyze investment in NbS and bird-friendly infrastructure to enhance the resilience of the Americas’ flyways by protecting, restoring and managing sites needed to secure at least 10% of the flyway populations of focal migratory species (CAF, 2024). Through a Green-Blue financing mechanism, AFI will initially support about 30 projects over the next decade, combining CAF funding with international cooperation, philanthropy, and private investment. Initial projects include, for instance, coastal climate-resilience actions in the Rocuant–Andalién wetland in Chile (CAF, 2024), a site identified in the shorebird minimum area portfolio with a high importance score (Figure 2A).

Our framework further supports AFI’s adaptive management approach by recognizing the dynamic nature of migratory bird populations and conservation networks. Population estimates and habitat conditions change over time, restoration actions can improve population status, and the securing of individual sites alters network irreplaceability. Accordingly, we recommend updating this prioritization as sites are added to the AFI portfolio, or, at minimum, revisiting it on a biennial basis to incorporate new data on species, populations, and conservation designations (e.g., Plumptre et al., 2025). Additionally, the workflow developed in this study can be applied by proponents outside the current priority areas to assess and demonstrate their eligibility for investment (Figure S1).

While our results offer a robust framework for informing fund allocation, some limitations must be acknowledged. Data availability remains uneven across regions and species. Crowd-sourced information such as eBird is geographically biased toward countries with long-standing monitoring traditions, particularly the United States (La Sorte & Somveille, 2020). In contrast, tropical regions, where many temperate-breeding migrants spend most of the year, lack information on populations and migration patterns (Bayly et al., 2018). This gap is even larger for austral migrants which move entirely within South America (Jahn et al., 2013, 2017). Expanding monitoring for these under-sampled areas and species will be crucial for refining future iterations of this analysis. Additionally, expanding the availability and accessibility of tracking data would facilitate the explicit integration of migratory connectivity in our analysis, enabling evaluation of each site’s role in maintaining functional connectivity within the portfolio (Dhanjal-Adams et al., 2017; Lin et al., 2020b). Finally, our threat assessment was limited to pressures that can be mapped at hemispheric scales (Seavy et al., 2025), and we evaluated them as presence/area rather than estimating population-level effects. Consequently, our results should be viewed as a scoping assessment of prevalent and extensive threats across spatial planning units, and site-level project design should be refined with local data and expertise (DeLuca et al., 2023).

Despite these limitations, our analysis provides a transparent and reproducible foundation for hemispheric conservation planning. We identify spatial priorities for migratory birds across the Americas using a full annual cycle approach that is explicitly designed to guide AFI’s implementation. The complementary shorebird and landbird portfolios bridge the gap between broad-scale (DeLuca et al., 2023) and site-level prioritizations, providing AFI with a practical blueprint for action. Integrating these spatial priorities with scalable NbS and adaptive monitoring can support targeted investment, help halt migratory bird declines, and strengthen the long-term resilience and biodiversity recovery of the Americas’ flyways.

## Acknowledgments

We are grateful to the teams and partner organizations that contributed data for this analysis, including the eBird Status and Trends project, the Atlantic, Midcontinental, and Pacific Shorebird Conservation Initiatives, Manomet, BirdLife International’s World Bird and Biodiversity Database, and the many local teams that proposed IBAs and WHSRN sites across the Americas. We also acknowledge the community scientists and field observers whose contributions to eBird, GBIF, tracking data (DeLuca et al. 2023, Table S3), and regional monitoring programs made this work possible. Finally, we thank Stuart Butchart and Martín Brasa for their helpful comments that improved this manuscript.

