## Supplementary information for "Operationalizing site-level conservation for migratory birds across the Americas’ flyways"

**Supplementary File 1.** Shorebird population data sources from literature.

Agreda, A. E. 2017. Plan de Conservación para Aves Playeras en Ecuador. Resumen Ejecutivo. Aves y Conservación / BirdLife en Ecuador, Red Hemisférica de Reservas para Aves Playeras. Quito, Ecuador. Pp. 58.

Andres, B. A., Smith, P. A., Morrison, R. I. G., Gratto-Trevor, C. L., Brown, S. C., & Friis, C. A. (2012). Population estimates of North American shorebirds, 2012.

Clay, R. P., Lesterhuis, A. J., & Johnson O. 2010. Conservation Plan for the American GoldenPlover (Pluvialis dominica). Version 1.1. Manomet Center for Conservation Sciences, Manomet, Massachusetts, USA.

Clay, R. P., Lesterhuis, A. J., & Centrón, S. 2012. Conservation Plan for the Lesser Yellowlegs (Tringa flavipes). Version 1.0. Manomet Center for Conservation Sciences, Manomet, Massachusetts, USA.

Clay, R. P., Lesterhuis A. J., Schulte, S., Brown, S., Reynolds, D., & Simons T. R. 2010. Conservation Plan for the American Oystercatcher (Haematopus palliatus) throughout the Western Hemisphere. Version 1.1. Manomet Center for Conservation Sciences, Manomet, Massachusetts, USA.

Fellows, S. D., & Jones S. L. 2009. Status assessment and conservation action plan for the Long-billed Curlew (Numenius americanus). U.S. Department of Interior, Fish and Wildlife Service, Biological Technical Publication, FWS/BTP-R6012-2009, Washington, D.C, USA.

Fernández, G., Buchanan, J. B., Gill, R. E., Lanctot, R. Jr., & Warnock, N. 2010. Conservation Plan for Dunlin with Breeding Populations in North America (Calidris alpina arcticola, C. a. pacifica, and C. a. hudsonia). Version 1.1. Manomet Center for Conservation Sciences, Manomet, Massachusetts, USA.

Fernández, G., Warnock, N., Lank, D. B., & Buchanan, J. B. 2010. Conservation Plan for the Western Sandpiper (Calidris mauri). Version 1.1. Manomet Center for Conservation Sciences, Manomet, Massachusetts, USA.

García, W. J., Senner, N., Norambuena, H., & Schmitt, F. (2018). Atlas de las aves playeras de Chile: sitios importantes para su conservación.

Lanctot, R. B., Aldabe, J., Almeida, J. B., Blanco, D., Isacch J. P., Jorgensen, J., Norland, S., Rocca P., & Strum K M. 2010. Conservation Plan for the Buff-breasted Sandpiper (Tryngites subruficollis). Version 1.1. U. S. Fish and Wildlife Service, Anchorage, Alaska, and Manomet Center for Conservation Sciences, Manomet, Massachusetts, USA.

Lesterhuis, A. J., & Clay, R. P. 2010. Conservation Plan for Wilson’s Phalarope (Phalaropus tricolor). Version 1.1. Manomet Center for Conservation Sciences, Manomet, Massachusetts, USA.

Payne, L. X. 2010. Conservation Plan for the Sanderling (Calidris alba). Version 1.1. Manomet Center for Conservation Sciences, Manomet, Massachusetts, USA.

Senner, N. R. 2010. Conservation Plan for the Hudsonian Godwit. Version 1.1. Manomet Center for Conservation Science, Manomet, Massachusetts, USA.

Senner, N. R. & Angulo, F. (2014). Atlas de las Aves Playeras del Perú: Sitios Importantes para su Conservación.

Wilke, A. L., & Johnston-González, R. 2010. Conservation Plan for the Whimbrel (Numenius phaeopus). Version 1.1. Manomet Center for Conservation Sciences, Manomet, Massachusetts, USA.

Zdravkovic, M. G. 2013. Conservation Plan for the Wilson’s Plover (Charadrius wilsonia). Version 1.0. Manomet Center for Conservation Sciences, Manomet, Massachusetts, USA

**Table S1.** Final list of focal shorebird (n = 51) and landbird (n = 61) species selected for analysis.

| Group | Scientific name | Common name |
| --- | --- | --- |
| Shorebirds | Recurvirostra americana | American Avocet |
| Shorebirds | Pluvialis dominica | American Golden-Plover |
| Shorebirds | Haematopus palliatus | American Oystercatcher |
| Shorebirds | Scolopax minor | American Woodcock |
| Shorebirds | Calidris bairdii | Baird's Sandpiper |
| Shorebirds | Arenaria melanocephala | Black Turnstone |
| Shorebirds | Pluvialis squatarola | Black-bellied Plover |
| Shorebirds | Himantopus mexicanus | Black-necked Stilt |
| Shorebirds | Calidris subruficollis | Buff-breasted Sandpiper |
| Shorebirds | Calidris alpina | Dunlin |
| Shorebirds | Thinocorus orbignyianus | Gray-breasted Seedsnipe |
| Shorebirds | Tringa melanoleuca | Greater Yellowlegs |
| Shorebirds | Limosa haemastica | Hudsonian Godwit |
| Shorebirds | Charadrius vociferus | Killdeer |
| Shorebirds | Calidris minutilla | Least Sandpiper |
| Shorebirds | Tringa flavipes | Lesser Yellowlegs |
| Shorebirds | Numenius americanus | Long-billed Curlew |
| Shorebirds | Limnodromus scolopaceus | Long-billed Dowitcher |
| Shorebirds | Haematopus leucopodus | Magellanic Oystercatcher |
| Shorebirds | Pluvianellus socialis | Magellanic Plover |
| Shorebirds | Limosa fedoa | Marbled Godwit |
| Shorebirds | Charadrius montanus | Mountain Plover |
| Shorebirds | Calidris melanotos | Pectoral Sandpiper |
| Shorebirds | Charadrius melodus | Piping Plover |
| Shorebirds | Calidris maritima | Purple Sandpiper |
| Shorebirds | Calidris canutus | Red Knot |
| Shorebirds | Phalaropus fulicarius | Red Phalarope |
| Shorebirds | Phalaropus lobatus | Red-necked Phalarope |
| Shorebirds | Calidris ptilocnemis | Rock Sandpiper |
| Shorebirds | Arenaria interpres | Ruddy Turnstone |
| Shorebirds | Charadrius modestus | Rufous-chested Dotterel |
| Shorebirds | Calidris alba | Sanderling |
| Shorebirds | Charadrius semipalmatus | Semipalmated Plover |
| Shorebirds | Calidris pusilla | Semipalmated Sandpiper |
| Shorebirds | Limnodromus griseus | Short-billed Dowitcher |
| Shorebirds | Charadrius nivosus | Snowy Plover |
| Shorebirds | Tringa solitaria | Solitary Sandpiper |
| Shorebirds | Actitis macularius | Spotted Sandpiper |
| Shorebirds | Calidris himantopus | Stilt Sandpiper |
| Shorebirds | Calidris virgata | Surfbird |
| Shorebirds | Oreopholus ruficollis | Tawny-throated Dotterel |
| Shorebirds | Charadrius falklandicus | Two-banded Plover |
| Shorebirds | Bartramia longicauda | Upland Sandpiper |
| Shorebirds | Tringa incana | Wandering Tattler |
| Shorebirds | Calidris mauri | Western Sandpiper |
| Shorebirds | Numenius phaeopus | Whimbrel |
| Shorebirds | Calidris fuscicollis | White-rumped Sandpiper |
| Shorebirds | Tringa semipalmata | Willet |
| Shorebirds | Steganopus tricolor | Wilson's Phalarope |
| Shorebirds | Charadrius wilsonia | Wilson's Plover |
| Shorebirds | Gallinago delicata | Wilson's Snipe |
| Landbirds | Buteo swainsoni | Swainson's Hawk |
| Landbirds | Petrochelidon pyrrhonota | Cliff Swallow |
| Landbirds | Sporophila ruficollis | Dark-throated Seedeater |
| Landbirds | Elanoides forficatus | Swallow-tailed Kite |
| Landbirds | Sporophila cinnamomea | Chestnut seedeater |
| Landbirds | Sporophila palustris | Marsh seedeater |
| Landbirds | Dolichonyx oryzivorus | Bobolink |
| Landbirds | Chordeiles minor | Common Nighthawk |
| Landbirds | Coccyzus americanus | Yellow-billed Cuckoo |
| Landbirds | Progne subis | Purple Martin |
| Landbirds | Tyrannus tyrannus | Eastern Kingbird |
| Landbirds | Contopus sordidulus | Western Wood-pewee |
| Landbirds | Contopus cooperi | Olive-sided Flycatcher |
| Landbirds | Chaetura pelagica | Chimney Swift |
| Landbirds | Cardellina canadensis | Canada warbler |
| Landbirds | Setophaga cerulea | Cerulean Warbler |
| Landbirds | Setophaga striata | Blackpoll Warbler |
| Landbirds | Coccyzus erythropthalmus | Black-billed Cuckoo |
| Landbirds | Pheucticus ludovicianus | Rose-breasted Grosbeak |
| Landbirds | Geothlypis philadelphia | Mourning Warbler |
| Landbirds | Empidonax traillii | Willow Flycatcher |
| Landbirds | Setophaga pensylvanica | Chestnut-sided Warbler |
| Landbirds | Protonotaria citrea | Prothonotary Warbler |
| Landbirds | Vermivora chrysoptera | Golden-winged Warbler |
| Landbirds | Icterus spurius | Orchard Oriole |
| Landbirds | Icterus galbula | Baltimore Oriole |
| Landbirds | Geothlypis formosa | Kentucky Warbler |
| Landbirds | Parkesia motacilla | Louisiana Waterthrush |
| Landbirds | Passerina cyanea | Indigo Bunting |
| Landbirds | Antrostomus carolinensis | Chuck-will's-widow |
| Landbirds | Archilochus colubris | Ruby-Throated Hummingbird |
| Landbirds | Empidonax minimus | Least Flycatcher |
| Landbirds | Helmitheros vermivorum | Worm-eating Warbler |
| Landbirds | Hylocichla mustelina | Wood Thrush |
| Landbirds | Passerina caerulea | Blue Grosbeak |
| Landbirds | Setophaga coronata | Yellow-rumped Warbler |
| Landbirds | Cardellina pusilla | Wilson's Warbler |
| Landbirds | Setophaga discolor | Prairie Warbler |
| Landbirds | Setophaga chrysoparia | Golden-cheeked Warbler |
| Landbirds | Vireo bellii | Bell's Vireo |
| Landbirds | Limnothlypis swainsonii | Swainson's Warbler |
| Landbirds | Calothorax lucifer | Lucifer Hummingbird |
| Landbirds | Leiothlypis virginiae | Virginia's Warbler |
| Landbirds | Selasphorus rufus | Rufous Hummingbird |
| Landbirds | Leiothlypis crissalis | Colima Warbler |
| Landbirds | Selasphorus calliope | Calliope Hummingbird |
| Landbirds | Vireo atricapilla | Black-capped Vireo |
| Landbirds | Catharus bicknelli | Bicknell's Thrush |
| Landbirds | Oporornis agilis | Connecticut Warbler |
| Landbirds | Calamospiza melanocorys | Lark Bunting |
| Landbirds | Micrathene whitneyi | Elf Owl |
| Landbirds | Anthus spragueii | Sprague's Pipit |
| Landbirds | Calcarius ornatus | Chestnut-collared Longspur |
| Landbirds | Antrostomus vociferus | Eastern Whip-poor-will |
| Landbirds | Selasphorus sasin | Allen's Hummingbird |
| Landbirds | Rhynchophanes mccownii | Thick-billed Longspur |
| Landbirds | Ammospiza leconteii | LeConte's Sparrow |
| Landbirds | Centronyx henslowii | Henslow's sparrow |
| Landbirds | Zonotrichia querula | Harris's Sparrow |
| Landbirds | Psiloscops flammeolus | Flammulated Owl |
| Landbirds | Setophaga kirtlandii | Kirtland's Warbler |

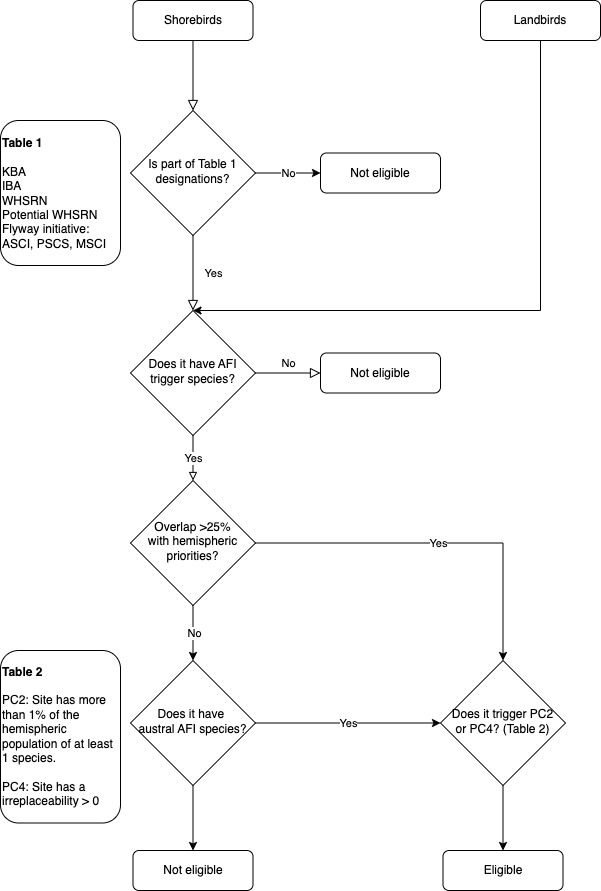

**Figure S1.** Decision tree outlining the stepwise criteria to determine eligibility of spatial planning units for AFI investment.
